# Channel Capacity Computations for Unregulated and Autoregulated Gene Expression

**DOI:** 10.1101/802108

**Authors:** Zahra Vahdat, Karol Nienałtowski, Zia Farooq, Michał Komorowski, Abhyudai Singh

## Abstract

How living cells can reliably process biochemical cues in the presence of molecular noise is not fully understood. Here we investigate the fidelity of information transfer in the expression of a single gene. We use the established model of gene expression to examine how precisely the protein levels can be controlled by two distinct mechanisms: (i) the transcription rate of the gene, or (ii) the translation rate for the corresponding mRNA. The fidelity of gene expression is quantified with the information-theoretic notion of information capacity. Derived information capacity formulae reveal that transcriptional control generally provides a tangibly higher capacity as compared to the translational control. We next introduce negative feedback regulation in gene expression, where the protein directly inhibits its own transcription. While negative feedback reduces noise in the level of the protein for a given input signal, it also decreases the input-to-output sensitivity. Our results show that the combined effect of these two opposing forces is a reduced capacity in the presence of feedback. In summary, our analysis presents the first analytical quantification of information transfer in simple gene expression models, which provides insight into the fidelity of basic gene expression control mechanisms.

## I. INTRODUCTION

Within the noisy environment of a living cell, biochemical species such as genes, RNAs, and proteins often occur at low molecular counts. These low copy numbers, coupled with the inherent random nature of biochemical processes, drive considerable stochastic fluctuations in copy numbers over time. Advances in single-cell technologies over the last decade have precisely quantified fluctuations in the level of a protein within single cells in diverse organisms [1]–[6]. There is also increasing appreciation of the beneficial and harmful roles played by these fluctuations with tangible consequences for biology and medicine [7]–[13].

As in engineering systems, cells use regulatory mechanisms to maintain the level of a protein around a set point. The simplest example of this is negative autoregulation, where a protein expressed from its gene inhibits its own transcription creating a feedback loop [14]–[23]. Such feedbacks have been shown to be ubiquitous across genes [24], and design of synthetic feedbacks for controlling gene expression is an intense area of current research [25]–[27]. In this contribution, we use information theory to study precision with which different mechanisms can control gene expression levels. Specifically, we deploy the concept of information capacity that has recently been used to study information processing in diverse living systems [28]–[32]. Information capacity is expressed in bits, and an input-output system with the capacity of *c* bits can reliably discriminate 2^*c*^ input states from the output. Here, the main focus is to quantify the information capacity of the gene expression process for the input considered to be at different locations of the gene expression process, and also when the expression operates in open loop vs. feedback loop.

We modify a well-studied model of stochastic protein synthesis from its corresponding gene with the inclusion of an input signal, where the input either enhances the transcription rate of the gene, or the translation rate for the corresponding mRNA (Fig. 1). Analytical formulas for the channel capacity are derived as a function of the fold change in the input from its basal to its maximum value. While it is not surprising that the channel capacity monotonically increases with the input fold change, the location of the input leads to contrasting results. More specifically, input-driven transcription generally provides a much higher capacity as compared to input-driven translation, but we also identify parameter regimes where the opposite is true. We also consider the case of autoregulated gene expression where the protein inhibits its own transcription. Interestingly, the inclusion of a negative feedback reduces the capacity irrespective of where the input affects gene expression.

**Fig. 1.**
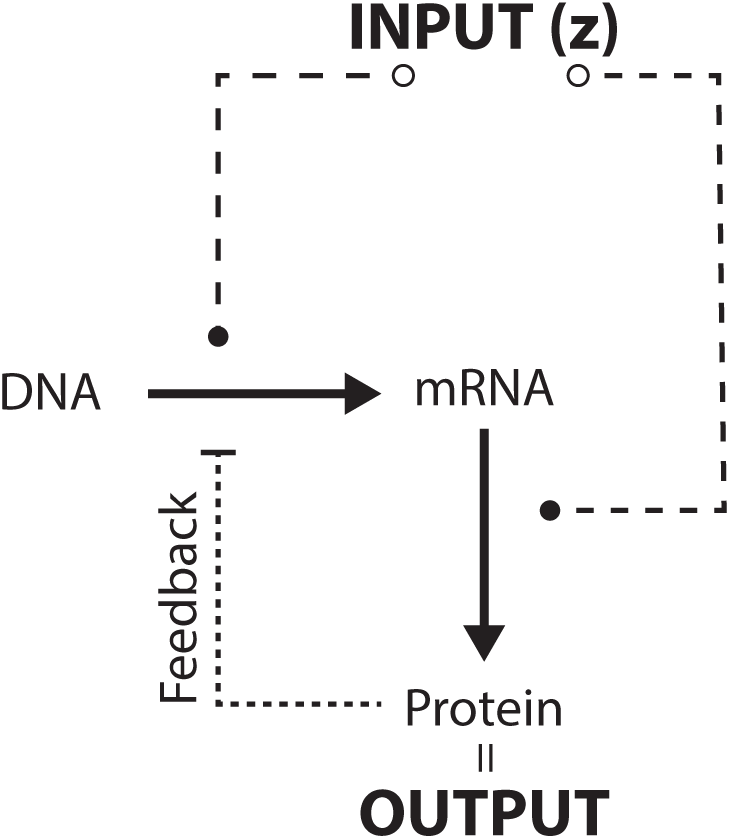
Schematic of the process of gene expression where the input signal may control either transcription or translation. Further, protein may inhibit its own transcription, which constitutes the negative feedback. The protein level is considered as the output of the system. We examine how precisely the input signal may control the output through transcription or translation, and with or without feedback.

The paper is organized as follows. In Section II we formulate a stochastic model of gene expression without feedback, and quantify fluctuations in the output protein level for a given input signal. In Section III we derive formulas for the channel capacity and systematically compare scenarios where the input affects transcription vs. translation. Negative feedback is introduced in Section IV with capacity comparisons to the no feedback case. Finally, conclusions are presented in Section V.

## II. STOCHASTIC MODELS OF GENE EXPRESSION

Motivated by experimental data on gene activity in individual cells, the simplest mechanistic model of gene expression considers a protein being synthesized in stochastic bursts [33]–[42]. The burst events are assumed to arrive as per a Poisson process with rate (burst frequency) *a*. Each burst event leads to a several proteins being made instantaneously. Let **x**(*t*) denote the concentration of a given protein within a cell at time *t*. The burst size, i.e., the size of the jump in **x**(*t*), is an independent and identically distributed random variable following an exponential distribution with mean burst size *b* [43], [44]. Assuming that the protein has a long half-life with no active degradation, then in between two successive burst events the protein level is diluted as per

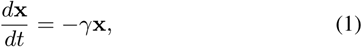

where *γ* is the cell growth rate. In essence, time evolution of **x**(*t*) is a Piecewise-Deterministic Markov Process (PDMP) with continuous exponential decay, interspersed with random burst events that increase protein levels at discrete times (as determined by the Poisson arrival process). Physiologically, the burst frequency *a* corresponding to the rate at which a gene is transcribed to produce mRNAs. Since mRNAs have short half-lives compared to the protein, each mRNA transcript quickly degrades after producing a burst of proteins [45], [46]. In this context, the burst size is the number of proteins made in the lifetime of an individual mRNA, and the average burst size *b* is directly related to the rate at which the mRNA is translated.

Having defined a PDMP model for bursty gene expression, the probability density function of the protein level at time *t, f* (*x, t*), evolves as per the following continuous master equation

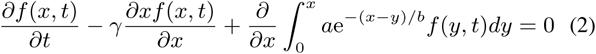

[47]–[49]. Prior work has elegantly shown that solving this master equation at equilibrium (*∂f* (*x, t*)*/∂t* = 0), yields the steady-state distribution for the protein level

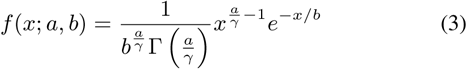

to be a gamma distribution, where Γ is the gamma function [47]. In the remainder of the paper, we set *γ* = 1 and *a* can be considered as the burst frequency normalized by the cell growth rate. The steady-state statistical moments of the protein level are

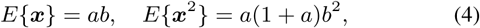

where *E* denotes the expected value operation evaluated at steady-state. Interestingly, experimental measurements have found several proteins in *E. coli* and budding yeast to show statistical fluctuations consistent with a gamma distribution [43], [44].

In order to compute the channel capacity, we next consider an input signal *z* that activates gene expression in two different ways:

1. The input increases the transcription rate of the gene, and this corresponds to a monotonically increasing burst frequency *a*(*z*) with respect to the input.
2. The input increases the translation rate of the mRNA, and this corresponds to a monotonically increasing average burst size *b*(*z*) with respect to the input.

For simplicity, we assume a Michaelis–Menten form for these functions

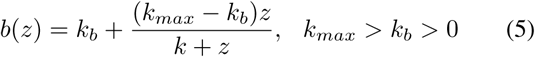

where *k*_*b*_ is the basal rate when there is no input (*z* = 0), *k*_*max*_ is the maximum rate (*z* → ∞). The constant *k* represents the value of the input when the burst frequency reaches the halfway point, i.e., *b*(*z*) = (*k*_*max*_ + *k*_*b*_)*/*2 when *z* = *k*, and thus *k* is inversely related to how rapidly *b*(*z*) increases with increasing *z* to reach *k*_*max*_. An identical functional form is assumed for *a*(*z*). Having quantified statistical fluctuations in the output protein level as a function of the input signal, we next focus our attention on the channel capacity.

## III. CHANNEL CAPACITY FOR UNREGULATED GENE EXPRESSION

Channel capacity has been proposed initially to quantify the information transfer over a noisy communication channel [50]. The transfer was assumed to take place between the input, i.e., sender’s end, and the output, i.e., the receiver’s end. Due to noise, the input cannot be reconstructed from the output without error. Information capacity quantifies the overall fidelity of the transmission. Broadly speaking, it quantifies the *log*_2_ of the maximal number of different messages that can be sent over the channel, on average, per transmission. Here, we use the capacity to quantify the fidelity of gene expression control. We assume that an input signal impacts the output protein level via the burst size or the burst frequency. The capacity between the input and output quantities, therefore, is the fidelity of the control mechanism in terms of the overall number of distinct input values that can be discriminated from the output. The exact computation of the information capacity is usually problematic. Therefore, we use the approximation given by the asymptotic capacity. The asymptotic capacity calculates the information capacity exactly for a large number of statistically independent copies of a given signaling system and then regresses back the capacity of a single copy [51].

Asymptotic capacity *C*_*A*_ is computed from the Fisher information matrix (FIM). For illustration, lets take an arbitrary probability distribution *f* with 2 parameters (*θ*_*i*_, *i* = 1, 2). Then the FIM is a 2 × 2 matrix with elements

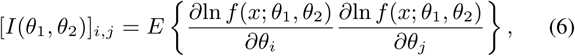

where *E* stands for expected value. Considering parameters *θ*_*i*_ depend on input signal *z, z* ∈ [0, *z*_*max*_], then the Fisher information for signal *z* is

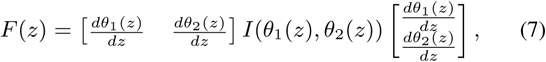

and the general formulation for asymptotic channel capacity can be computed from the Fisher information using

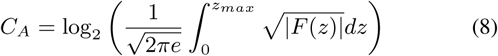

[51]. For the gamma distribution derived in (3), the Fisher information matrix is given by

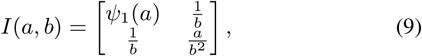

where *ψ*_1_(.) is Trigamma function (the logarithmic second derivative of gamma function),

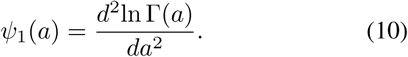

Next we study specific cases when either *a* or *b* is modulated as a Michaelis–Menten function of the input *z*.

### A. Input activates gene expression via the burst size

In this case the mean burst size varies with *z* as

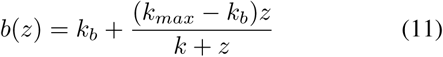

and the burst frequency *a* is constant. Using (7) and (9) we calculate the Fisher information as

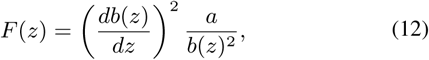

that simplifies to

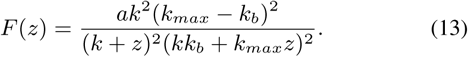

Finally, we calculate the asymptotic channel capacity from (8)

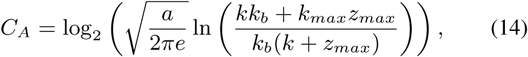

which in the limit (*z*_*max*_ → ∞) reduces to

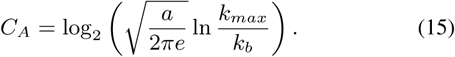

### B. Input activates gene expression via the burst frequency

In this case the burst frequency varies with *z* as

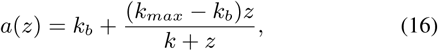

and the mean burst size *b* is constant. Using (7), (9) and (16) yields the following Fisher information

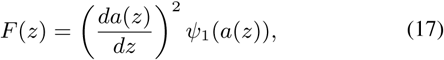

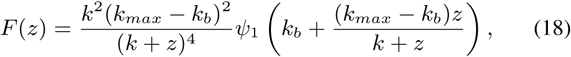

with the asymptotic channel capacity from (8) as

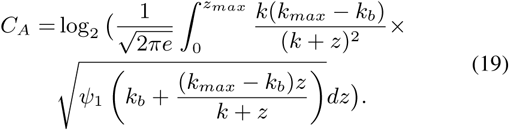

The integration within (19) can be further simplified by doing a change of variable

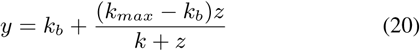

which in the limit *z*_*max*_ → ∞ simplifies (19) to

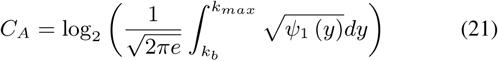

with *ψ*_1_ being the Trigamma function defined in (10).

The channel capacities derived in (15) and (21) provide some intriguing insights:

- The capacities only depends on *k*_*max*_ and *k*_*b*_, the maximum and minimum value of the input, respectively, and are invariant of the parameter *k* in (16) and (11).
- When the input drives the burst size *b*, then the capacity (15) monotonically increases with the burst frequency as in 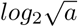.
- When the input drives the burst frequency *a*, then the capacity (19) becomes independent of the average burst size *b*, and hence, independent of the expression specific parameters.
- As illustrated in Fig. 2, for sufficiently small *a* the input in burst frequency (i.e., transcription) has higher capacity than input in burst size (i.e., translation) irrespective of *k*_*max*_ and *k*_*b*_.
- For sufficiently large *a*, the input in burst size can provide higher capacity provided the ratio *k*_*max*_*/k*_*b*_ is small enough (Fig. 2).

## IV. CHANNEL CAPACITY FOR AUTOREGULATED GENE EXPRESSION

Negative feedback is incorporated in the gene expression process by modifying the burst frequency to be a monotonically decreasing function of the protein level **x**

**Fig. 2.**
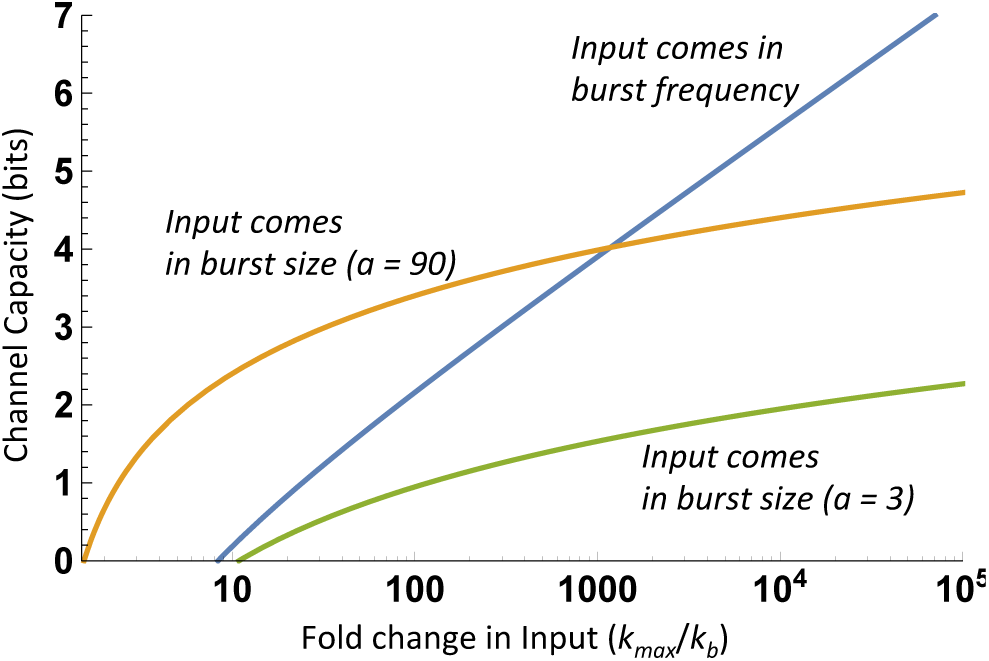
Channel capacity for unregulated stochastic gene expression when the input activates transcription via the burst frequency, or activates translation via the burst size. Channel capacities derived in (15) and (21) are plotted as function of *k*_*max*_ for different value of *a* assuming *k*_*b*_ = 1. The input affecting transcription generally provides a higher capacity. However, our results show that when *a* is large and *k*_*max*_ is small, then input-driven translation can provide a higher capacity.

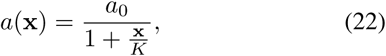

with two new positive parameters *a*_0_ and *K*. Here *a*_0_ is the burst frequency when there is no feedback (*K* → ∞), and 1*/K* can be intuitively thought of as the feedback strength, i.e., lower values of *K* correspond to stronger feedback. In this case, the steady-state distribution of the protein concentration is given by

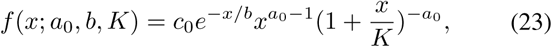

[52], where *c*_0_ is the normalization constant

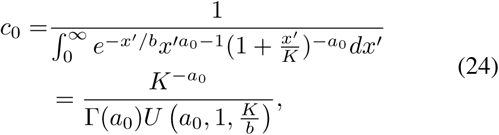

and *U* is the confluent hypergeometric function. This distribution yields the following steady-state first and second order moments

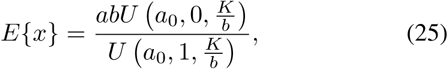

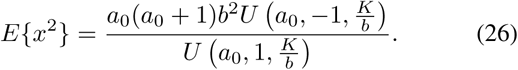

As before, we now consider two different cases of input-dependent expression, but now with negative feedback.

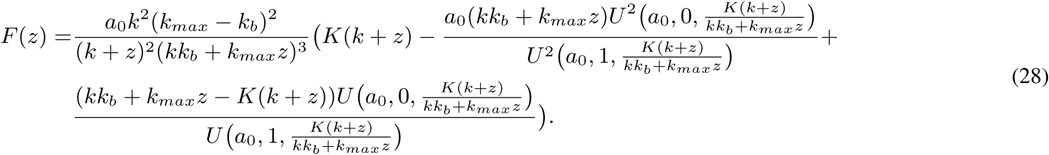

### A. Input activates gene expression via the burst size

Considering the input activates gene expression by enhancing mRNA translation, we model the average burst size as a function of input *z* as per (11), while the burst frequency is given by (22). Following (6), (7), and (23) the Fisher information *F* (*z*) is calculated in Appendix A and is given by (28). The channel capacity (*C*_*A*_) is computed via (8) using (28). Given the complexity of the formula we investigate it numerically. Our results show that decreasing values of *K*, which corresponds to increasing negative feedback strength, results in a reduced capacity (Fig. 3).

**Fig. 3.**
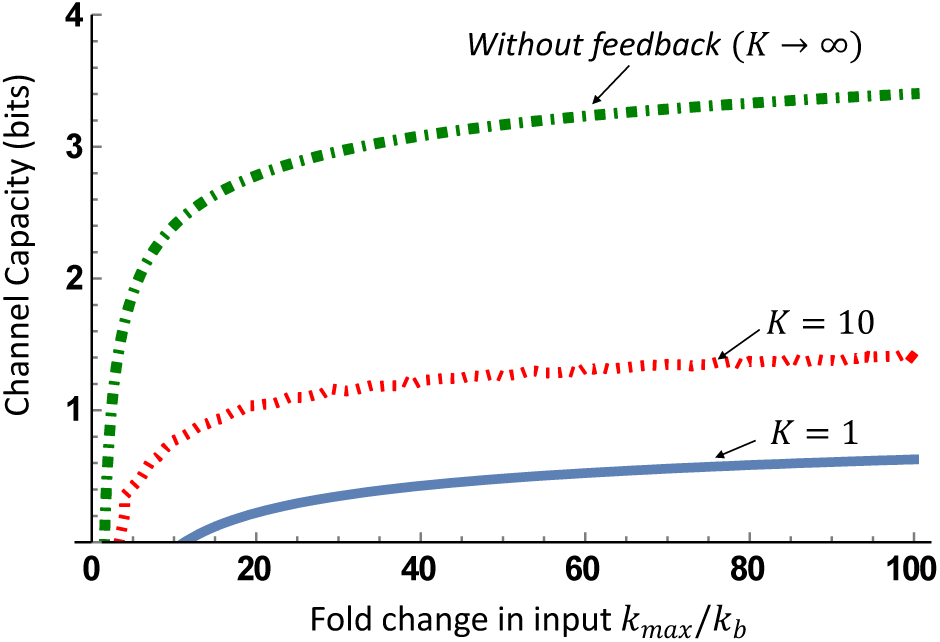
Channel capacity for negative feedback regulation of burst frequency, and the input affects the mean burst size. The asymptotic channel capacity (*C*_*A*_) is computed via (8) using (28) for parameters *a*_0_ = 90, *k*_*b*_ = 1, *k* = 1. Decreasing values of *K* (i.e., increasing negative feedback strength) results in a reduced information capacity.

### B. Input activates gene expression via the burst frequency

Next, we consider the input signal affecting the burst frequency. Since both the input and the feedback come at the same location, we modify the burst frequency to

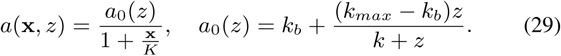

The mean burst size is now assumed to be a constant *b*. The Fisher information for input signal *z* is given by (30) (see Appendix B for details), where *U*_(1,0,0)_ and *U*_(2,0,0)_ are the first and second derivative of the confluent hypergeometric function with respect to *a*_0_(*z*), respectively. Calculating *C*_*A*_ from (8) again shows that adding negative feedback reduces the information capacity (Fig. 4).

**Fig. 4.**
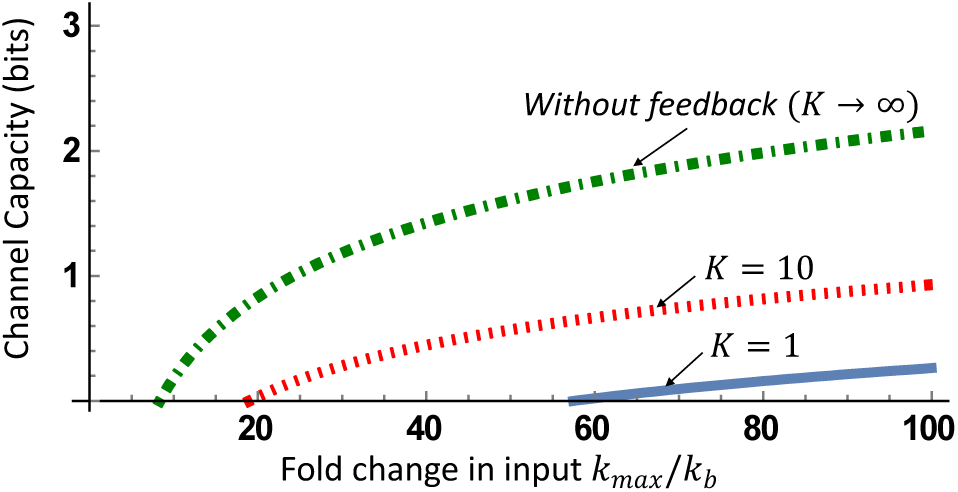
Channel capacity for negative feedback regulation of burst frequency, and the input also affects the burst frequency. The asymptotic channel capacity (*C*_*A*_) is computed via (8) using (30) for parameters *b* = 5, *k*_*b*_ = 1, *k* = 1. As in Fig. 3, decreasing values of *K* (i.e., increasing negative feedback strength) result in a reduced information capacity.

## V. CONCLUSION

We have systematically investigated fidelity of information transfer in a simple yet mechanistic model of stochastic gene expression. In the absence of any feedback loops, our results provide simple analytical formulas connecting the information capacity to properties of the input signal (the maximum and minimum values of the input), and expression parameters. A key finding from this analysis is that transcriptional control of gene expression generally leads to a higher capacity as compared to translational control (Fig. 2). However, if the burst frequency is high enough, then translational control can provide a higher capacity for small fold changes in the input (Fig. 2).

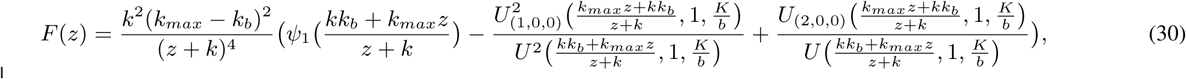

These results were extended to consider feedback in burst frequency. In the feedback case, the formulas are quite involved and we have investigated them numerically. It is quite possible that formulas can be further simplified in limits of strong (*K* → 0) or weak feedback (*K* → ∞) and we will investigate this is our future work. Our results lead to an intriguing find that in spite of feedback loops suppressing noise in protein levels for a given input value [16], [53], they actually reduce the information capacities (Figs. 3 and 4). Intuitively, inclusion of feedback makes the output less sensitive to changes in the input, and the overall effect of noise buffering and reduced input-to-output sensitivity is a lower information capacity. While our analysis is restricted to a simple non-cooperative feedback, it will be interesting to see if more complex forms of feedback can lead to a higher capacity.

Another direction of future work is to consider feedbacks in the burst size and/or the protein decay rate. Our recent collaborative work has derived probability density functions for such control mechanisms [49], and we plan to investigate channel capacities across feedback architectures. Prior work has shown the negative feedback is much more efficient in suppressing external disturbances in protein production, rather than noise arising from the inherent bursty nature of expression [16]. We plan to expand models presented here to incorporate external disturbances by making parameters itself random processes that biologically corresponds to fluctuations in the abundance of enzymes involved in gene expression. With multiple noise mechanism affecting the protein level, we will study information capacity both with and without feedback.

## ACKNOWLEDGMENT

AS is supported by the National Science Foundation grant ECCS-1711548 and ARO grant W911NF-19-1-0243. ZF, KN, and MK were supported by the Foundation for Polish Science within the First TEAM (First TEAM/2017-3/21) programme co-financed by the European Union under the European Regional Development Fund.

## VI. APPENDIX

### A. Calculating Fisher information when an input activates gene expression via the burst size

Fisher information is calculated by

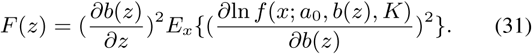

Same as (35) we have

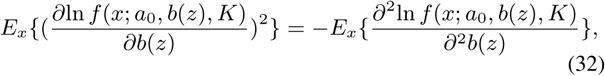

where

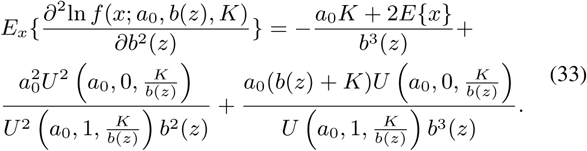

*E* {*x*} is calculated in (25). Finally using the above equations we get *F* (*z*) in (28).

### B. Calculating Fisher information when an input activates gene expression via the burst frequency

Since *a*_0_ is a function of input *z, a*_0_(*z*) changes as per (16), with constant burst size, following (7) Fisher information is as follows

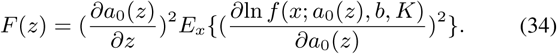

As 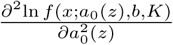 exists,

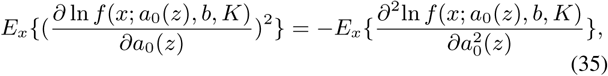

where

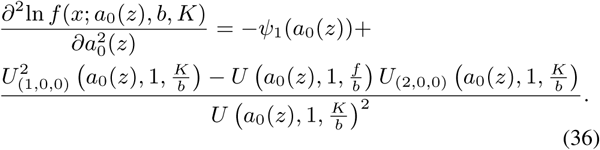

Finally, *F* (*z*) can be calculated by using (34), (35) and (36).

